# Oral octanoylcarnitine alleviates exercise intolerance in mouse models of long-chain fatty acid oxidation disorders

**DOI:** 10.1101/2025.06.24.661304

**Authors:** Keaton J. Solo, Yuxun Zhang, Sivakama S. Bharathi, Bob B. Zhang, Adam C. Richert, Alexandra V. Schmidt, Shakuntala Basu, Clinton Van’t Land, Olivia D’Annibale, Timothy C. Wood, Jerry Vockley, Eric S. Goetzman

## Abstract

Long-chain fatty acid oxidation disorders (LC-FAODs) cause energy deficits in heart and skeletal muscle that is only partially corrected by current medium-chain lipid therapies such as triheptanoin. We find that heart and muscle lack medium-chain acyl-CoA synthetases, limiting the capacity for β-oxidation of medium-chain fatty acids. Instead, heart and muscle mitochondria robustly respire on medium-chain acylcarnitines. The mitochondrial matrix enzyme carnitine acetyltransferase (CrAT) efficiently converts orally delivered octanoylcarnitine (C_8_-carnitine) to octanoyl-CoA for energy generation. C_8_-carnitine exhibits twice the oral bioavailability of triheptanoin and distributes to muscle and heart. A single oral dose markedly enhances grip strength, basal locomotion, and treadmill endurance while attenuating lactate and creatine kinase elevations in multiple mouse models of LC-FAODs. Thus, medium-chain acylcarnitines overcome a previously unrecognized metabolic bottleneck in LC-FAOD muscle and may represent an alternative to triglyceride-based therapies for bioenergetic disorders.

## INTRODUCTION

Long-chain fatty acid oxidation disorders (LC-FAODs) are a group of rare, inherited metabolic conditions caused by defects in mitochondrial enzymes required for the β-oxidation of long-chain fatty acids. There are two subgroups of LC-FAODs. The first group are defects in the carnitine shuttle, which facilitates transport of long-chain fatty acids into the mitochondria. This group includes mutations in the genes carnitine palmitoyltransferase-1a (CPT1a), carnitine-acylcarnitine translocase (CACT), and carnitine palmitoyltransferase-2 (CPT2). The second group are mutations in mitochondrial matrix β-oxidation enzymes, most notably very long-chain acyl-CoA dehydrogenase (VLCAD) and mitochondrial trifunctional protein (TFP). The two LC-FAOD subgroups exhibit largely overlapping symptoms due to impaired energy production, particularly during periods of fasting or increased energy demand [1, 2]. The major organs affected in patients during infancy are liver and heart. Depending on the severity of the mutation, mortality during this period can be high. Later, in adolescence, muscle dysfunction becomes the dominant symptom moving into adulthood. Most adult patients suffer recurring, life-threatening bouts of rhabdomyolysis, increasing the disease burden on both patients and healthcare systems [1, 3].

Current treatment for LC-FAODs consists of frequent feeding and replacement of dietary long-chain fats with medium-chain fats [4]. For 40+ years, medium-chain fats were provided as medium-chain triglycerides (MCT) containing C_8_ fatty acids. The dogma of MCT oil, which currently has a ∼$2.5 billion annual market as a supplement for weight management and sports nutrition, is that the C_8_ fatty acids traverse the mitochondrial membrane without the need for facilitated transport by the carnitine shuttle and rapidly produce energy via β-oxidation [5]. In LC-FAODs, MCT adequately managed liver symptoms but often failed to address cardiomyopathy and rhabdomyolysis [6]. This was believed to be caused by chronic depletion of TCA cycle intermediates [7]. The proposed solution was to replace the C_8_ fatty acids with C_7_ fatty acids, which, after two rounds of chain-shortening, produce a C_3_ remnant which is converted to succinate, thereby refilling the TCA cycle (anaplerosis) [7]. Consequently, the preferential treatment for LC-FAODs is now triheptanoin, a C_7_ triglyceride. Triheptanoin is believed to possess the rapid energy production properties of MCT with the additional benefit of anaplerosis [8]. Clinical trials and case studies demonstrate that triheptanoin improves symptoms for many LC-FAOD patients [9, 10]. However, there are limitations to the utility of triheptanoin. First, it is not efficacious against the cardiomyopathy seen in patients with severe, neonatal onset LC-FAODs, a subgroup that continues to exhibit high mortality [11]. Second, triheptanoin is only partially effective against the rhabdomyolysis that dominates the clinical picture for patients who live past childhood [10]. And third, the rate of treatment-related adverse events with triheptanoin is very high, ∼85%. Diarrhea, abdominal pain, and vomiting caused by triheptanoin oil reduce the quality of life for many patients [10].

New therapies are needed that focus on reducing the symptoms of rhabdomyolysis in LC-FAODs. In the present studies, we used LC-FAOD mouse models to interrogate the metabolism of MCT and triheptanoin in muscle and heart. Our work identifies two major barriers to the use of MCT or triheptanoin as a therapy for LC-FAOD muscle symptoms. First, the bulk of medium-chain lipids taken orally are consumed by the liver. Second, and most importantly, muscle and heart have a low capacity for activating medium-chain fatty acids to coenzyme A (CoA), which is a prerequisite step for fatty acid β-oxidation. In short, our data challenge the dogma of MCT and triheptanoin metabolism for organs outside the liver. We further show that there is an alternative route to convert medium-chain fatty acids to their acyl-CoA forms in muscle, which is by supplying them as acylcarnitine esters and leveraging carnitine acyltransferase enzymes to exchange the carnitine for CoA. Finally, we establish the efficacy of acylcarnitine therapy for treating exercise intolerance in mouse models of LC-FAODs.

## RESULTS

### Medium-chain triglycerides fail to improve exercise capacity in an FAOD mouse model

In mice, long-chain acyl-CoA dehydrogenase (LCAD) is the dominant acyl-CoA dehydrogenase family member, fulfilling the role that VLCAD plays in humans [12-14]. Correspondingly, LCAD knockout mice (LCADKO) model the symptoms of human VLCAD deficiency, including hypoketotic hypoglycemia and cardiomyopathy [14, 15]. Here, we established that LCADKO mice also exhibit multiple signs of muscle dysfunction, such as significantly reduced basal locomotion, reduced grip strength, and reduced latency to fall on the hanging wire test (Supplemental Fig 1). When exercise-naïve LCADKO mice are challenged with a run-to-exhaustion protocol, they run only one-third the distance of their wildtype counterparts (Fig 1A). Despite running a much shorter distance than wildtype mice, LCADKO mice display post-run lactic acidosis (Fig 1B). Pre-run blood glucose, pre-run blood lactate, and post-run blood glucose values are not different from wildtype (Supplemental Fig 2). These data suggest an increased reliance upon glycolysis for energy during exercise. We then tested whether medium-chain lipid therapies could improve exercise performance in the LCADKO mouse model. FAOD patients consume MCT oil or triheptanoin at ∼0.5 g/kg three to four times per day, with an overall target of ∼35% of total calories [10]. To mimic this dosing, LCADKO mice were gavaged with 0.5 mg/g of MCT oil or triheptanoin 20 minutes prior to a treadmill challenge. Neither compound improved running distance or post-run lactic acidosis (Fig 1A, B). We also tested the effect of chronic feeding by incorporating MCT and triheptanoin into pelleted mouse diets at 35% of total calories. LCADKO mice were fed these diets for 10 days and then challenged with acute exercise. Again, neither compound improved running distance nor ameliorated the post-run lactic acidosis (Fig 1A, B).

**Figure 1.**
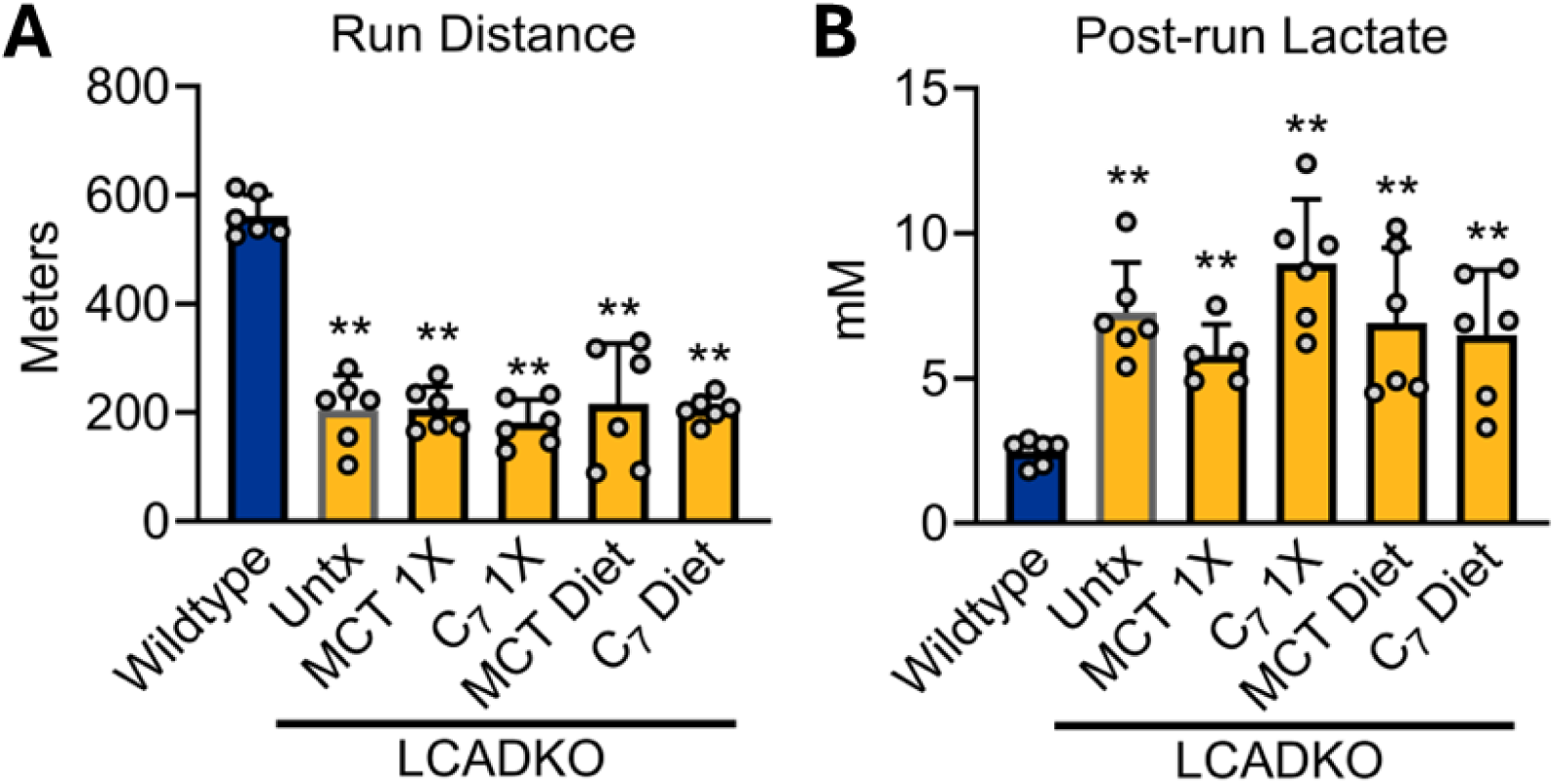
Medium-chain triglycerides fail to improve exercise capacity in an FAOD mouse model. Male LCADKO age 12-16 weeks, and matched wildtype controls, were run to exhaustion on a treadmill. Running distance (A) and blood lactate post-run (B) were recorded. Cohorts of LCADKO mice were either untreated (Untx) or treated with medium-chain triglycerides (MCT) or triheptanoin (C_7_). MCT and C_7_ were given either as 0.5 mg/g oral boluses (1X) or delivered as 35% of calories in pelleted diets for 10 days. None of the treatments improved running distance or lactic acidosis. **P value <0.01 for all LCADKO groups versus wildtype controls. All bars represent means and standard deviations.

### Exogenous medium-chain fatty acids are poorly metabolized outside of liver

We set out to determine why medium-chain therapies were inefficacious for treating exercise intolerance in the LCADKO mouse. First, we interrogated absorption and first-pass liver metabolism. It is generally held that oral medium-chain lipids are trafficked from the gut into portal circulation [16], but empirical data on the degree of first-pass liver metabolism of these lipids has been conflicting [17-19]. Further, little is known about the absorption of triheptanoin, which has supplanted MCT as the therapy of choice for FAODs. We interrogated the oral bioavailability of triheptanoin using rats with an indwelling jugular port. Rats were orally gavaged with triheptanoin or infused with the same amount of C_7_ fatty acids via the jugular port. Blood was sampled from the port at time points from 0 to 90 minutes. Blood lipids were hydrolyzed to convert any absorbed triheptanoin into free C_7_, which was then quantified by mass spectrometry. Oral bioavailability for triheptanoin was calculated as the ratio of area-under-the curve (AUC) for oral dosing to the AUC for i.v. dosing. Triheptanoin was observed to be ∼9% orally bioavailable (Fig 2A), meaning 9% of the C_7_ reaches systemic circulation. Thus, low bioavailability likely contributed to the lack of therapeutic effect observed in the LCADKO mouse model.

**Figure 2.**
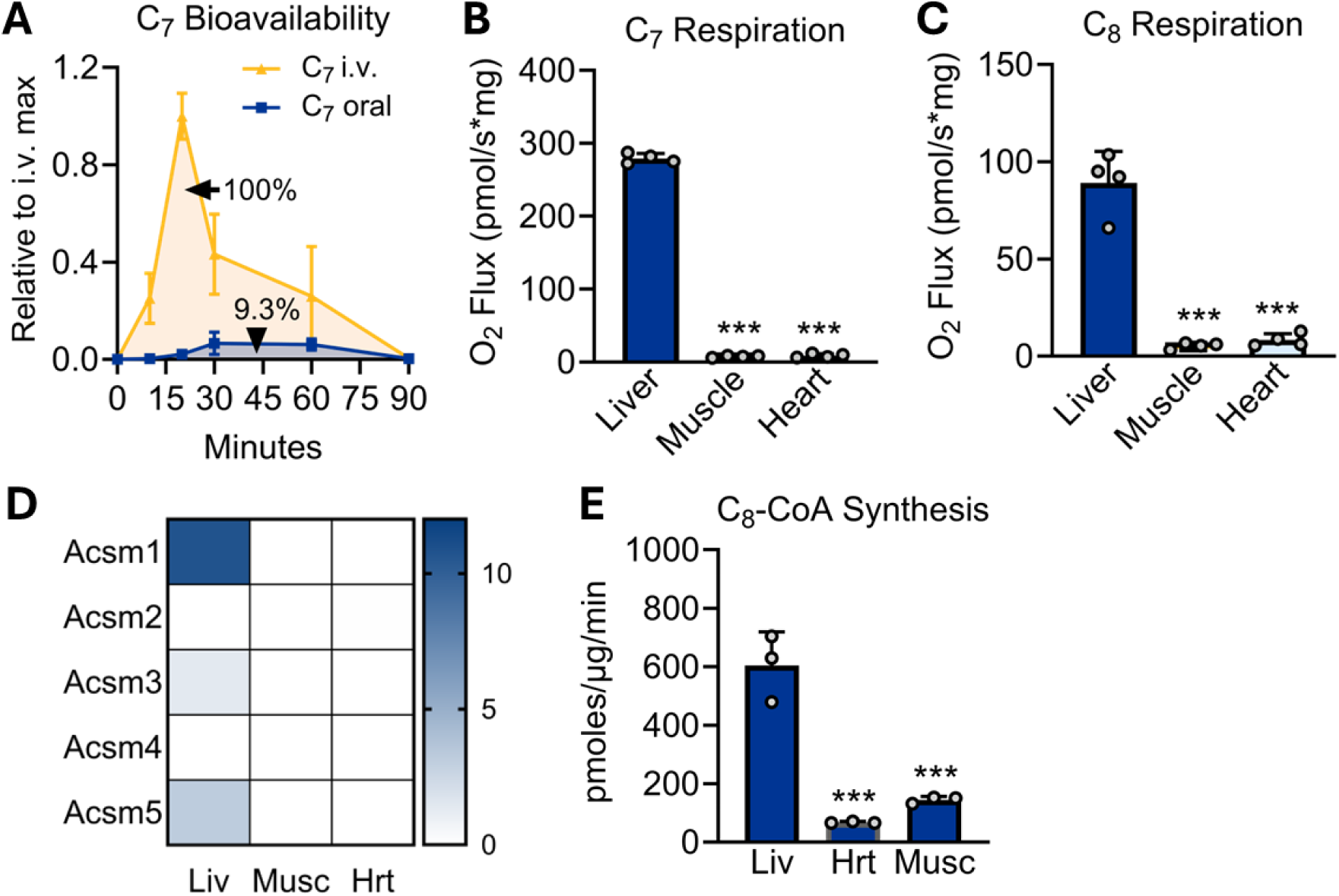
Exogenous medium-chain fatty acids are poorly metabolized outside of liver. A) Oral bioavailability of triheptanoin was determined by dosing N=3 male rats with 0.5 mg/g and following C_7_ fatty acid levels in the blood over time. The ratio of the area under the curve for oral dose: i.v. dose represents bioavailability, which is 9.3%. B,C) Mitochondria were isolated from the indicated wildtype mouse tissues and used for Oroboros respirometry studies to determine oxygen consumption on medium-chain free fatty acids C_7_ and C_8_. D) A previously published proteomics dataset [20] was interrogated for expression levels of the five medium-chain acyl-CoA synthetase (ACSM) isoforms. E) ^14^C-labeled C_8_ was followed to ^14^C_8_-CoA in the presence of mitochondria isolated from wildtype mouse liver, heart (Hrt), or muscle (Musc). ***P value <0.001, heart and muscle versus liver. All bars represent means and standard deviations.

We next asked whether, in the event systemic distribution could be improved, which medium-chain fatty acid would be preferred by the target organs muscle and heart. Oroboros mitochondrial respirometry was used to evaluate utilization of C_7_ and C_8_ by wildtype mouse muscle and heart mitochondria, with liver mitochondria serving as a benchmark control. Notably, both muscle and heart mitochondria exhibited very low capacities for oxidizing free C_7_ or free C_8_ fatty acids, as compared to liver (Fig 2B,C). To determine the mechanism behind this striking discrepancy we analyzed our recently published wildtype mouse proteomics dataset for expression of the medium-chain FAO machinery in liver, muscle, and heart [20]. All three tissues express the required enzymes for medium-chain FAO including medium-chain acyl-CoA dehydrogenase (MCAD), enoyl-CoA hydratase (ECHS1), hydroxyacyl-CoA dehydrogenase (HAD), and the medium-chain ketoacyl-CoA thiolase (THIM)(Supplemental Fig 3). However, muscle and heart completely lacked expression of the medium-chain acyl-CoA synthetases (ACSMs) which are required to activate medium-chain fatty acids to CoA for FAO (Fig 2D). There are five ACSM isoforms. ACSM2 is kidney-specific and ACSM4 is gut-specific [21-23]. Liver expresses the remaining three ACSMs (ACSM1, 3, and 5) but none of these were detected in muscle or heart. Immunoblotting corroborated these results for ACSM1 and ACSM3 (Supplemental Fig 4). Muscle, but not heart, exhibited potential low-level expression of ACSM5 (Supplemental Fig 4). In an ACSM activity assay, based on conversion of ^14^C-C_8_ into ^14^C-C_8_-CoA, wildtype muscle and heart homogenates exhibited 10-20% capacity for C_8_-CoA synthesis compared to liver homogenates (Fig 2E).

### Heart and muscle mitochondria prefer medium-chain acylcarnitines over medium-chain fatty acids

Pereyra et al [24] recently demonstrated that muscle and heart mitochondria have the capacity to oxidize medium-chain FA, but must do so via the carnitine shuttle, ala long-chain fatty acids. This is in keeping with our observation that muscle and heart lack ACSM expression. ACSMs are mitochondrial matrix enzymes. In liver, where ACSMs are abundantly expressed, medium-chain FA can flip-flop across the mitochondrial membranes and be activated to acyl-CoA in the matrix for FAO. In muscle and heart, the only path to FAO would be via long-chain acyl-CoA synthetases (ACSLs), which are known to have weak activity with medium-chain substrates [25, 26]. ACSLs are all extramitochondrial, or on the outside of the mitochondria facing the cytosol, and any medium-chain acyl-CoA formed by these enzymes would be forced to access the mitochondrial matrix through the carnitine shuttle. We therefore reasoned that muscle and heart mitochondria would be capable of oxidizing medium-chain acylcarnitines for energy. Indeed, both muscle and heart mitochondria robustly respired on C_7_-carnitine and C_8_-carnitine, although heart had a clear preference for C_8_-carnitine (Fig 3A,B). Such respiration would require the action of a carnitine acyltransferase enzyme, such as CPT2, to convert the metabolically inert medium-chain acylcarnitines into acyl-CoAs for oxidation. Wildtype mouse heart and muscle homogenates convert C_7_-carnitine to C_7_-CoA at a much faster rate than they convert free C_7_ to C_7_-CoA (Fig 3C,D). In contrast, liver shows no preference and equally converts C_7_ and C_7_-carnitine into C_7_-CoA (Fig 3E).

**Figure 3.**
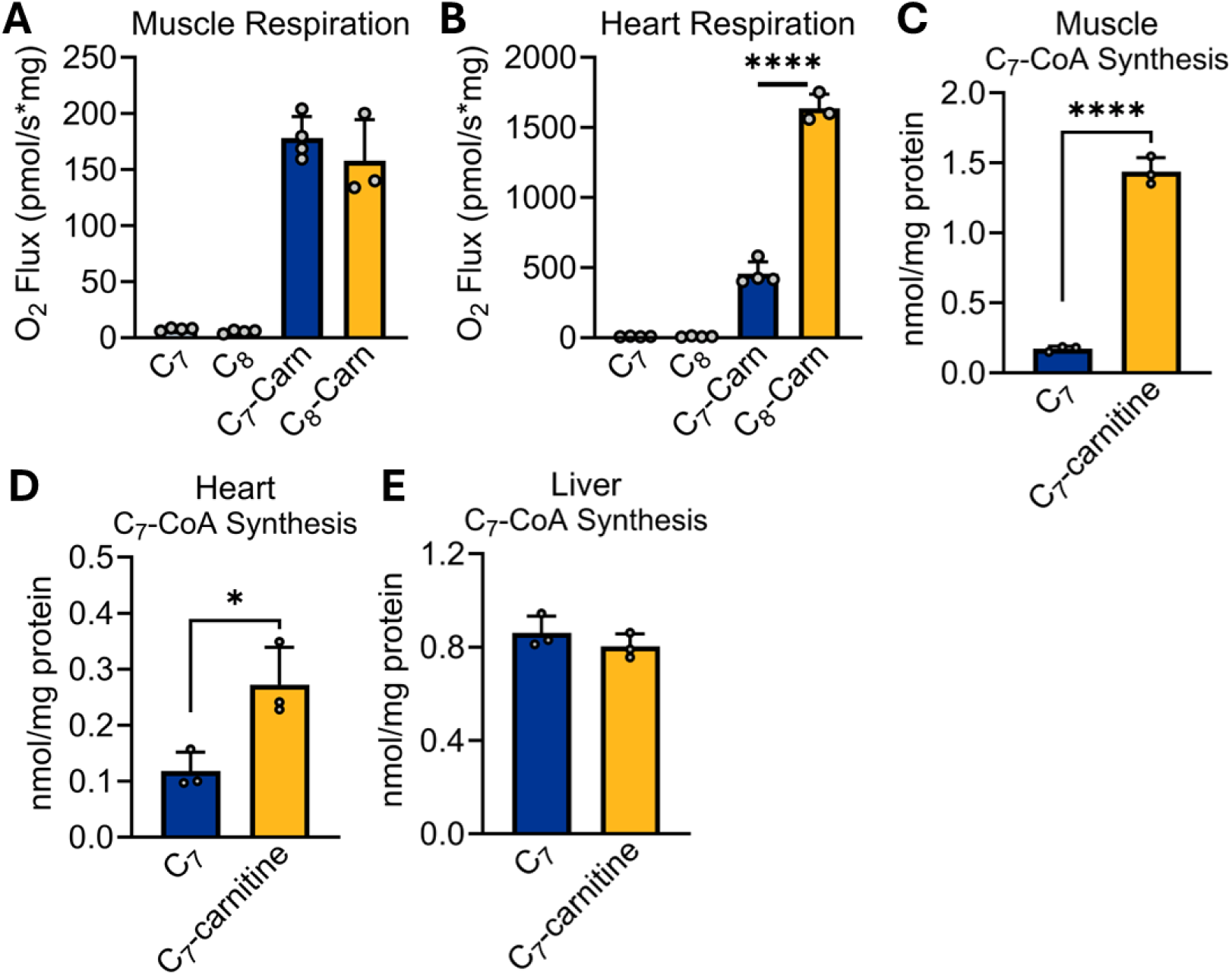
Heart and muscle mitochondria prefer medium-chain acylcarnitines over medium-chain fatty acids. A,B) Wildtype muscle and heart mitochondria were isolated and probed for respiratory capacity in the presence of either medium-chain free-fatty acids (C_7_, C_8_) or their acylcarnitine conjugates (C_7_-Carn, C_8_-Carn) using the Oroboros Oxygraph-2K. C-E) Wildtype mouse tissue homogenates were incubated with C_7_ or C_7_-carnitine. Mass spectrometry was used to quantify the amount of C_7_-CoA formed in one hour. *P<0.05; ****P<0.0001 by Student’s T-test. All bars represent means and standard deviations.

### Muscle and heart mitochondria use CrAT to metabolize medium-chain acylcarnitines

The carnitine shuttle canonically consists of CPT1, CACT, and CPT2 [2]. CPT1 converts acyl-CoAs to acylcarnitines at the outer membrane. The acylcarnitines are imported through the inner membrane by CACT, and then CPT2 reconverts them to acyl-CoAs for oxidation. We anticipated that CPT2 would be required for oxidation of medium-chain acylcarnitines by muscle and heart mitochondria. Unexpectedly, muscle from CPT2 muscle-specific conditional knockout mice (CPT2^mKO^) exhibited higher, not lower, C_7_-carnitine oxidation than floxed littermate controls (Fig 4A). The functional efficiency of the knockout was confirmed by a near-complete loss of mitochondrial respiration on the long-chain substrate palmitoylcarnitine (C_16_-carnitine). Further, in the acyl-CoA synthetase enzyme activity assay, muscle from CPT2^mKO^ mice showed a severely limited capacity to convert C_16_-carnitine to C_16_-CoA, while liver and heart from the same animals had a normal capacity for this reaction (Fig 4B). In agreement with the Oroboros respirometry data, muscle homogenates from CPT2^mKO^ mice converted C_7_-carnitine to C_7_-CoA at an accelerated rate compared to wildtype muscle (Fig 4C). Together, these data indicated that CPT2 is not required for mitochondria to oxidize exogenous medium-chain acylcarnitines. Heart and muscle are known to highly express a related enzyme, carnitine acetyltransferase (CrAT), which interconverts acetyl-CoA and acetylcarnitine [27]. Perhaps as a compensatory mechanism, CrAT expression is increased in CPT2^mKO^ muscle (Fig 4D). To test whether CrAT is responsible for mediating the oxidation of medium-chain acylcarnitines, we generated muscle-specific CrAT knockout mice (CrAT^mKO^). CrAT^mKO^ muscle mitochondria respired robustly on the CPT2 substrate C_16-_carnitine but could not respire on C_7_-carnitine (Fig 4E). CrAT’s optimal activity is with C_2_ substrates, but some evidence exists for activity with longer-chain substrates up to C_8_ [28, 29]. Using recombinant CrAT and CPT2, we confirmed that both enzymes can convert C_7_ and C_8_ acylcarnitines into their acyl-CoA counterparts *in vitro* (Fig 4F). The inability of CPT2 to facilitate medium-chain acylcarnitine oxidation *in vivo* suggests compartmentalization of medium-chain and long-chain acylcarnitine metabolism.

**Figure 4.**
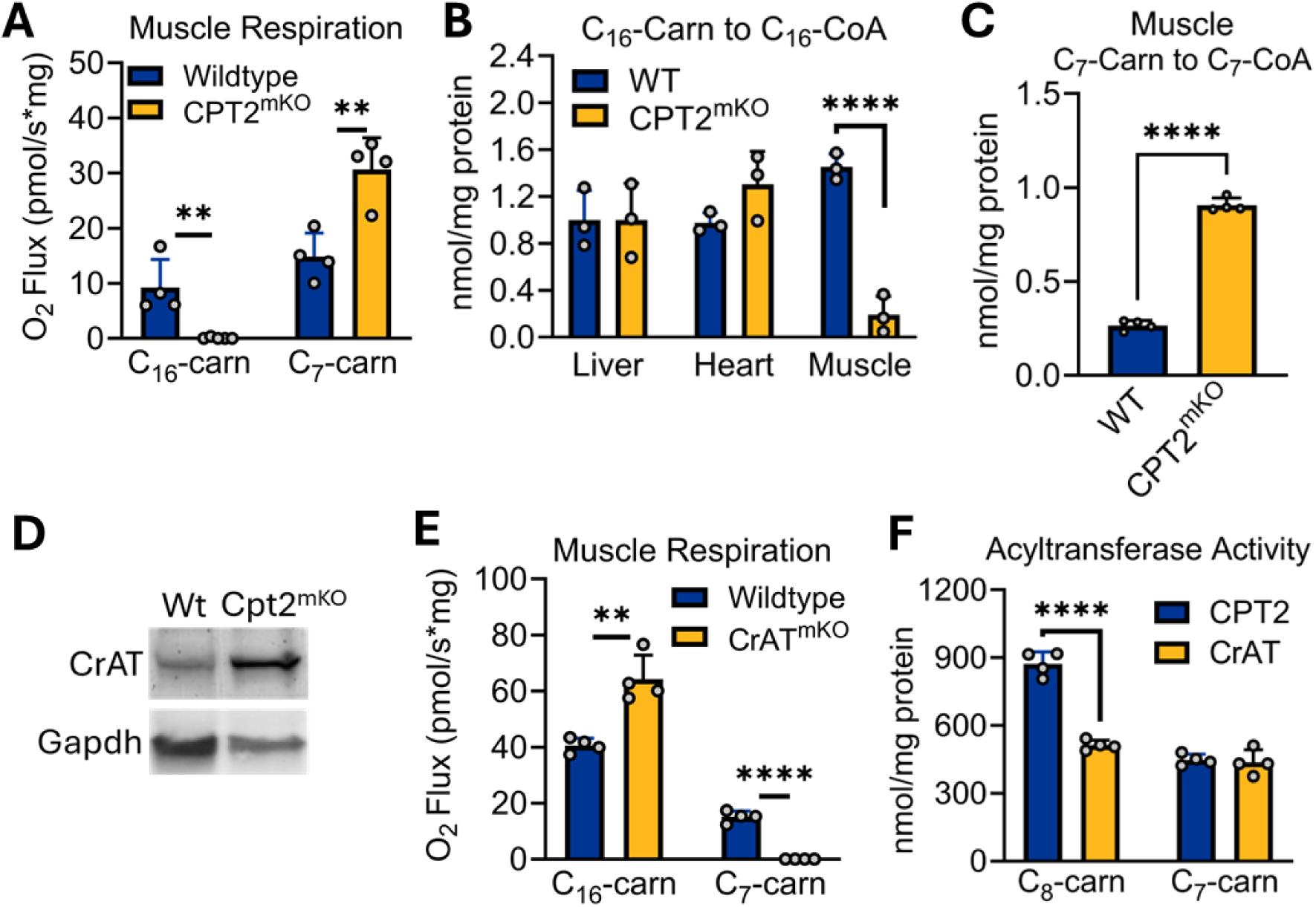
Muscle and heart mitochondria use CrAT to metabolize medium-chain acylcarnitines. A) Muscle mitochondria isolated from CPT2 muscle-specific knockout mice (CPT2^mKO^) cannot respire on C16-carnitine, the preferred substrate for CPT2, but respire robustly on C_7_-carnitine. B) The tissue specificity of the CPT2^mKO^ allele is demonstrated by the ability of liver and heart to normally convert C_16_-carnitine to C_16_-CoA, while muscle has near-zero capacity for this conversion. C) CPT2^mKO^ muscle mitochondria, while unable to convert C_16_-carnitine to C_16_-CoA (panel B), robustly convert C_7_-carnitine to C_7_-CoA. D) The significant increase in acyl-CoA synthetase activity in panel C is likely due to a compensatory increase in muscle CrAT in the CPT2^mKO^ mouse. E) Muscle mitochondria isolated from CrAT muscle-specific knockout mice (CrAT^mKO^) show the opposite result from CPT2^mKO^ muscle, i.e., high respiration on C_16_-carnitine but near-zero respiration on C_7_-carnitine. F) The intact mitochondria experiments shown in panels A and E indicate that CrAT is the major contributor to C_7_-carnitine conversion to C_7_-CoA, but *in vitro,* both CPT2 and CrAT are active with medium-chain acylcarnitine substrates. **P<0.01; ****P<0.0001 by Student’s T-test. All bars represent means and standard deviations.

### Oral C_8_-carnitine is rapidly absorbed, distributes to target organs, and improves exercise capacity in LCADKO mice

The studies above suggested the possibility that medium-chain acylcarnitines may have therapeutic value as an alternative energy source for FAODs. We therefore followed the fate of orally administered C_8_-carnitine. Using rats with jugular ports, as was done with triheptanoin in Fig 2A, we established that 18% of orally dosed C_8_-carnitine reaches system circulation intact (Fig 5A). Thus, C_8_-carnitine is about 2X more orally bioavailable than triheptanoin. Blood levels peaked at 10 minutes post-dosing, and the C_8_-carnitine was rapidly cleared. Similarly, when ^14^C-C_8_-carnitine tracer was administered orally, ^14^C-CO_2_ appeared in the exhaled breath within 10 minutes and metabolism was virtually complete after 1 hr (Fig 5B).

**Figure 5.**
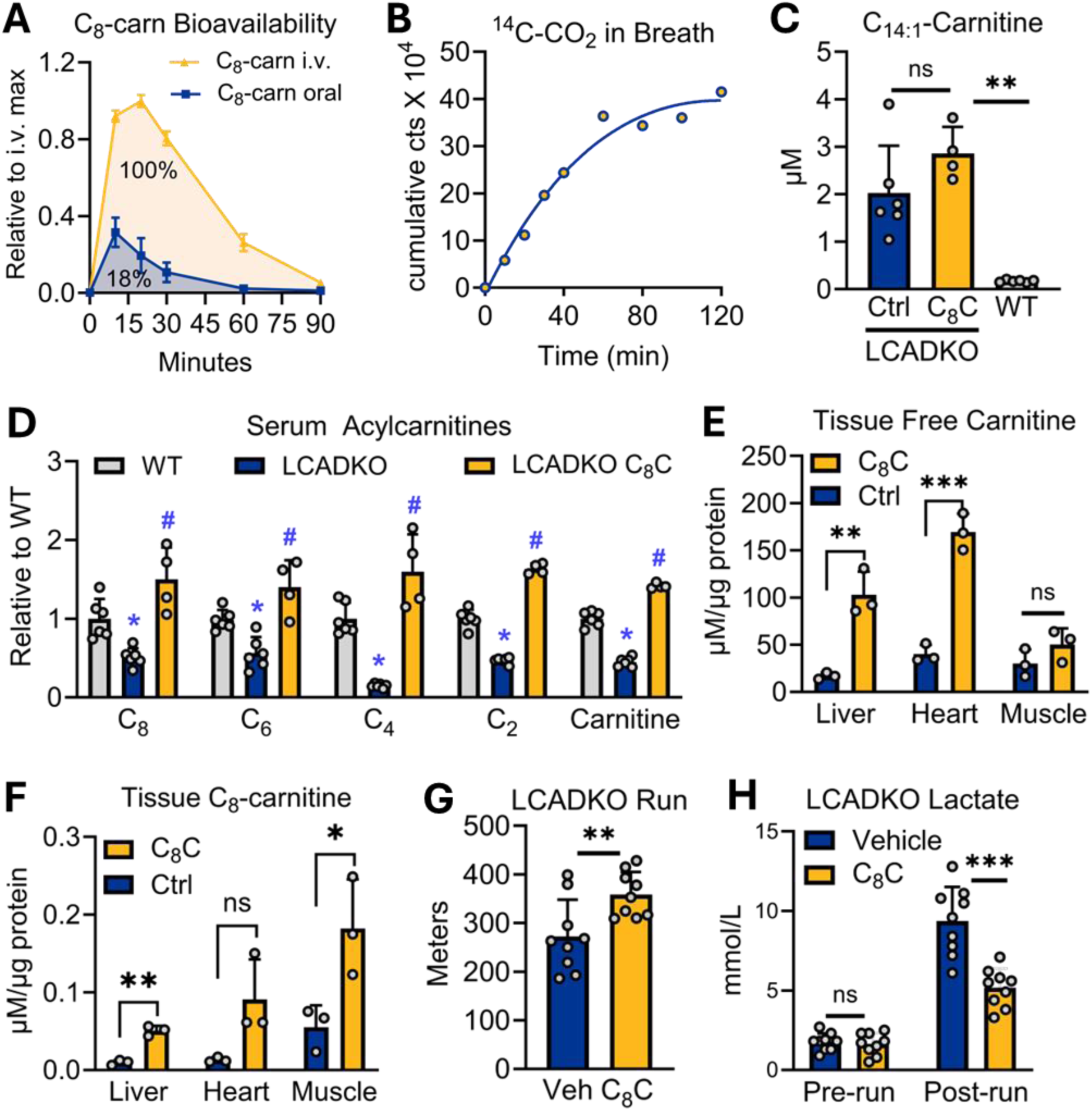
Oral C_8_-carnitine is rapidly absorbed, distributes to target organs, and improves exercise capacity in LCADKO mice. A) Oral bioavailability of C_8_-carnitine was determined by dosing N=3 male rats with 0.5 mg/g and following C_8_-carnitine levels in the blood over time. The ratio of the area under the curve for oral dose: i.v. dose represents bioavailability, which is 18%. B) Data are averages of two male wildtype mice gavaged with 3 µCi of ^14^C-labeled C_8_-carnitine. Mice were placed into boxes connected to a KOH trap to capture exhaled breath. Samples withdrawn from the trap at the indicated times were subjected to scintillation counting. The Y-axis is the cumulative counts over time. When the curve plateaus, it means the substrate has been completely oxidized. *C-F) Effect of feeding control diet (Ctrl) versus C_8_-carnitine (C_8_C) at 3% w/w for 10 days on serum and tissue acylcarnitines.* C) Serum C_14:1_-carnitine is pathognomonic for human VLCAD deficiency and is recapitulated in the LCADKO mouse. Dietary C_8_C does not alter this diagnostic long-chain acylcarnitine species. D) Serum acylcarnitines <8 carbons are systemically reduced in LCADKO mice (blue vs. gray bars); dietary C_8_C increases all to normal or above normal levels (yellow bars). E,F) Tissue levels of free carnitine and C_8_-carnitine are increased in the disease-relevant tissues liver, muscle, and heart after feeding C_8_-carnitine. G,H) Single gavaged doses of 0.5 mg/g C_8_-carnitine improve running capacity in LCADKO mice while limiting lactic acidosis. Panel B, *P<0.05 LCADKO versus wildtype and #P<0.05 LCADKO C_8_C versus wildtype; all other panels, **P<0.01; ***P<0.001 by Student’s T-test. All bars represent means and standard deviations.

Next, LCADKO mice were fed with either standard diet (Ctrl) or standard diet supplemented with 3% w/w C_8_-carnitine for 10 days. Serum, muscle, heart, and liver tissue were subjected to acylcarnitine profiling. Because of compromised ability to chain-shorten long-chain fatty acids, FAOD patients and mouse models accumulate long-chain acyl-CoAs inside the mitochondria but have a paucity of partially-shortened short/medium-chain CoAs. This pattern is reflected in the serum acylcarnitine profile and is diagnostic for these disorders [30]. Increased serum C_14:1_-carnitine is pathognomonic for VLCAD deficiency and is seen in LCADKO mice as a model of VLCAD deficiency [14, 15]. As expected, 10 days of dietary C_8_-carnitine did not significantly alter serum C_14:1_-carnitine levels (Fig 5C). However, dietary C_8_-carnitine increased the levels of downstream short-chain acylcarnitines, including free carnitine, all of which are normally suppressed in FAODs (Fig 5D). Tissue free carnitine and C_8_-carnitine levels were also significantly increased (Fig 5E,F). Together these data show that oral C_8_-carnitine distributes to key organs affected in FAODs, where its conversion to C_8_-CoAs produces energy but also boosts free carnitine levels, which can relieve mitochondrial stress in FAODs by promoting acylcarnitine efflux [4]. The therapeutic potential of C_8_-carnitine was evidenced by gavaging LCADKO mice with 0.5 mg/g C_8_-carnitine just prior to an acute exercise challenge, which significantly improved performance while also limiting lactic acidosis (Fig 5G,H).

### Oral C_8_-carnitine improves exercise capacity in a VLCAD-deficient mouse model

In mouse muscle, LCAD contributes ∼70% of the long-chain acyl-CoA dehydrogenase enzyme activity and VLCAD ∼30% [14, 15]. In humans, LCAD is not expressed in muscle, and therefore VLCAD contributes 100% [31]. Recently, a more clinically relevant model of VLCAD deficiency was developed by conditionally knocking out LCAD in skeletal muscle of a VLCAD^-/-^ mouse strain [32]. This animal demonstrates all the primary symptoms of human VLCAD deficiency, including hypoglycemia, cardiomyopathy, and myopathy with recurrent rhabdomyolysis. Here, this double knockout strain (VLCAD^dKO^) was used to assess the preclinical efficacy of oral C_8_-carnitine, compared to triheptanoin or vehicle (saline) as a control. A cohort of wildtype mice was included in these studies as a benchmark of normal function. First, we gavaged VLCAD^dKO^ mice with either C_8_-carnitine or triheptanoin at 0.5 mg/g (or saline as control), waited 20 minutes, and then monitored basal locomotion in an open-field apparatus. C_8_-carnitine treated VLCAD^dKO^ mice moved ∼2X more than either vehicle or triheptanoin-treated VLCAD^dKO^ mice, achieving the same locomotion as the wildtype group (Fig 6A). The amount of time spent resting and the speed of locomotion were also completely normalized by C_8_-carnitine, but not triheptanoin (Fig 6B,C). A single dose of C_8_-carnitine, but not triheptanoin, similarly normalized grip strength in the VLCAD^dKO^ mice to that of wildtype mice (Fig 6D). Further, in an acute exercise challenge, a single dose of C_8_-carnitine increased running distance by 5-fold in VLCAD^dKO^ mice, although this distance was still significantly shorter than that run by wildtype mice (Fig 6E). Despite running five-fold further, VLCAD^dKO^ mice did not develop the lactic acidosis seen in either vehicle or triheptanoin-treated VLCAD^dKO^ mice (Fig 6F). Finally, we observed that serum creatine kinase was significantly elevated in VLCAD^dKO^ mice even at baseline, without any exercise challenge (Fig 6G). Treadmill running increased serum CK by four-fold; gavaging with C_8_-carnitine significantly blunted this increase (Fig 6H). These data indicate that C_8_-carnitine has the potential to relieve muscle metabolic stress in FAODs.

**Figure 6.**
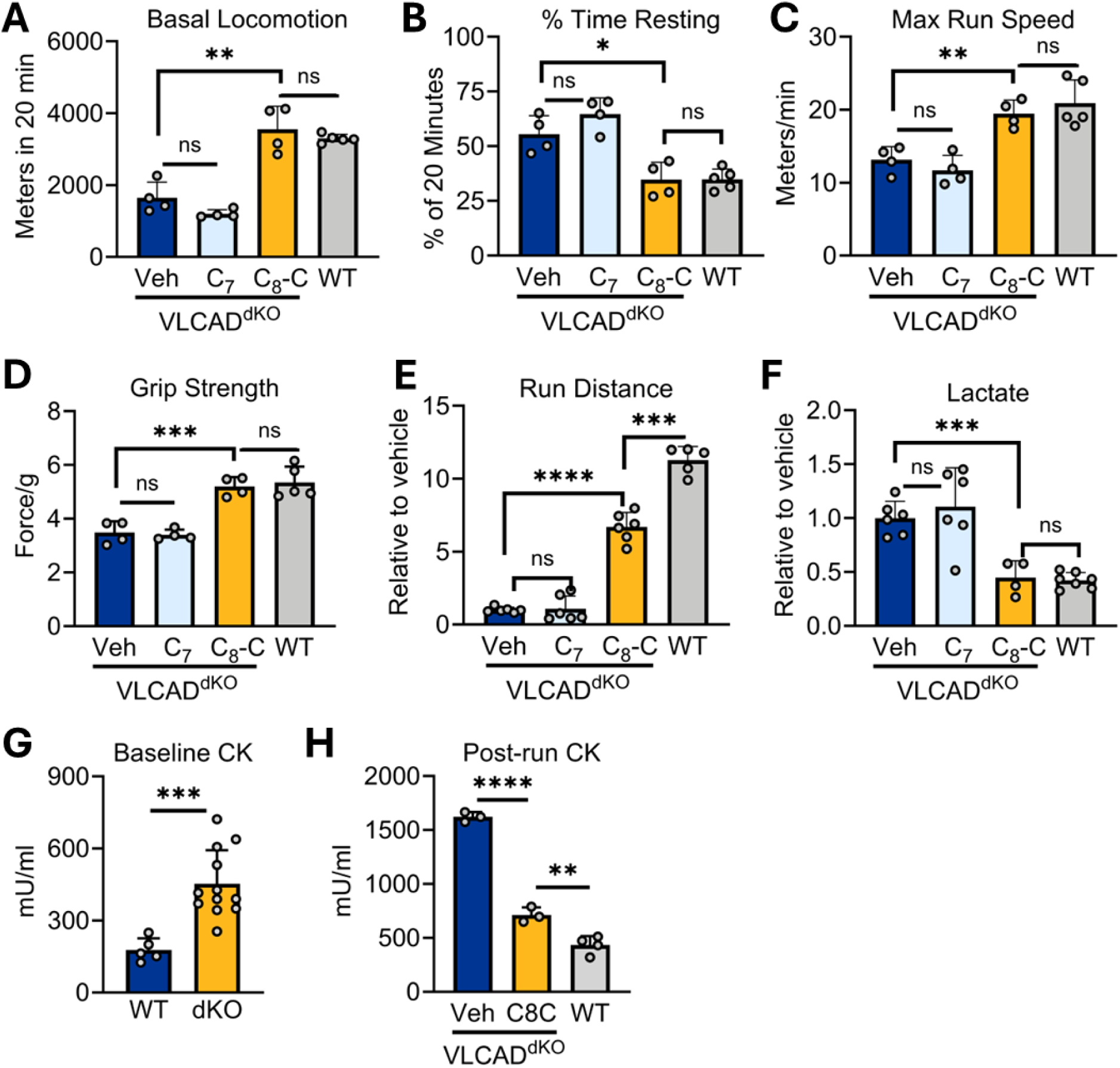
Oral C_8_-carnitine improves exercise capacity in a VLCAD-deficient mouse model. The blue and yellow bars in each graph represent groups of VLCAD-deficient mice that are double knockout (dKO) for LCAD in muscle tissue while gray bars are wildtype controls. A-C) An open-field actimeter was used to track basal locomotion behaviors in untreated VLCAD^dKO^ (Veh), triheptanoin-treated VLCAD^dKO^ mice (C_7_), or C_8_-carnitine treated VLCAD^dKO^ mice (C_8_C). C_7_ and C_8_C were given by gavage (0.5 mg/g) 20 min prior to placement in the open field apparatus. The apparatus measured total distance moved in 20 min (A), percent of time spent resting (B), and the maximal velocity reached while in motion (C). D) A force meter was used to measure forelimb grip strength, normalized to body weight. E,F) Mice were challenged with acute treadmill running to exhaustion. To combine data across runs, the data was normalized to untreated (Veh) VLCAD^dKO^ mice. Panel E is the distance run at exhaustion, panel F is post-run blood lactate. G,H) A separate cohort of mice was used to induce rhabdomyolysis by eccentric (downhill) running. After running to exhaustion, mice were returned to their home cages. 24 hr later blood was sampled for creatine kinase (CK) measurements. *P<0.05; **P<0.01; ***P<0.001; ****P<0.0001 by Student’s T-test. All bars represent means and standard deviations.

### Oral C_8_-carnitine improves muscle symptoms in CPT2-deficient, but not CrAT-deficient, mice

CPT2-deficient patients present with many of the same symptoms as VLCAD-deficient patients, including exercise-induced decompensation and rhabdomyolysis. Our data presented in Figure 4 indicate that CPT2 is not required for muscle C_8_-carnitine oxidation, and therefore, C_8_-carnitine may have utility for treating CPT2 deficiency. Complete ablation of CPT2 in mice is embryonic lethal. Muscle-specific knockouts exhibit much milder disease, characterized by compromised exercise performance [33]. Using the single-dose treatment paradigm studied above in LCADKO and VLCAD^dKO^ mice, we observed that C_8_-carnitine significantly increases running speed and grip strength in CPT2^mKO^ mice (Fig 7A,B). In contrast, muscle-specific CrAT^mKO^ mice did not show any improvement in these parameters after C_8_-carnitine dosing (Fig 7C,D). This strongly suggests that CrAT, and not CPT2, is required for the therapeutic benefit of medium-chain acylcarnitines for treating FAODs.

**Figure 7.**
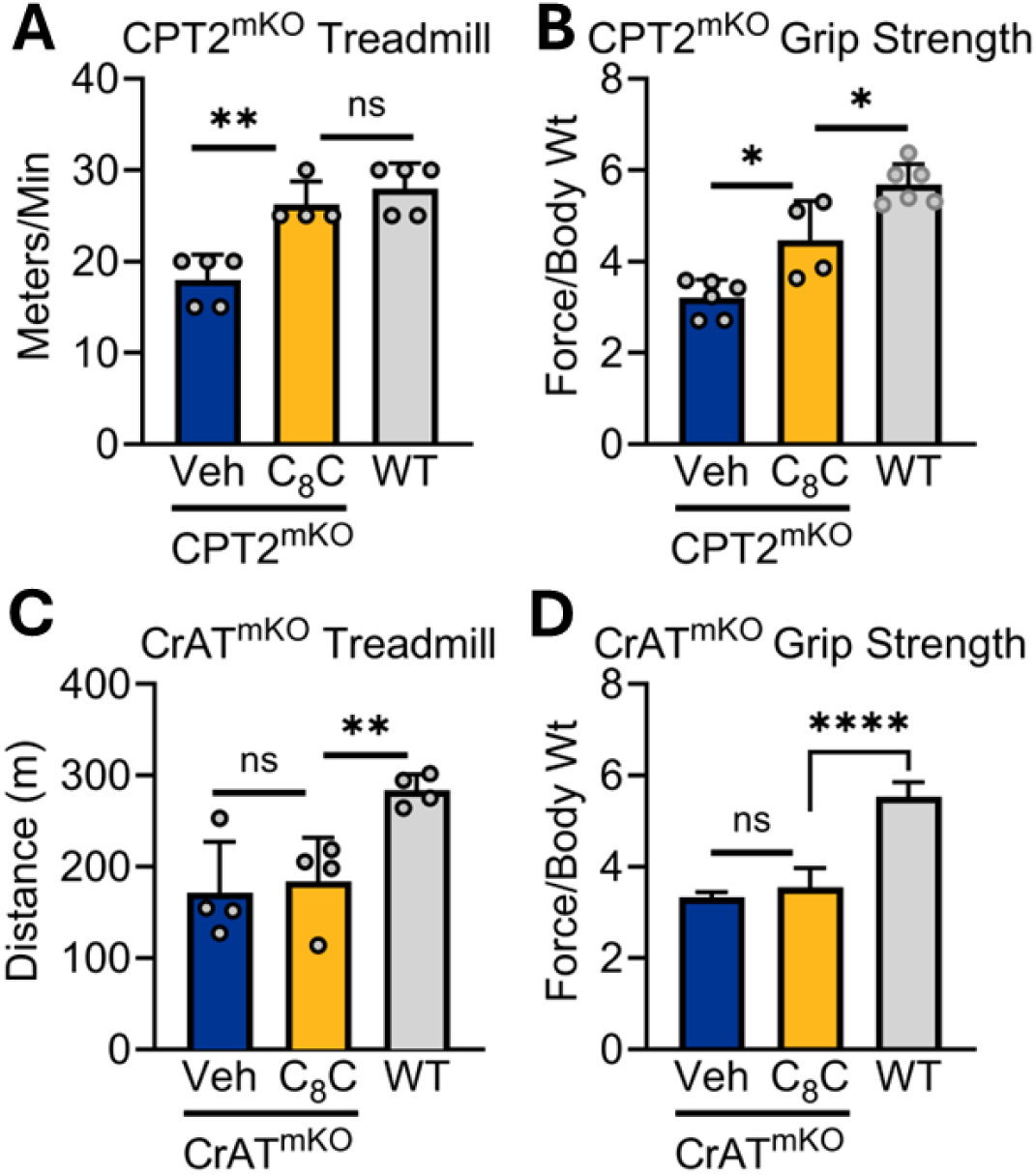
Oral C_8_-carnitine improves muscle symptoms in CPT2-deficient, but not CrAT-deficient, mice. A,B) Muscle-specific knockouts of CPT2 (CPT2^mKO^) were given saline vehicle (Veh) or C_8_-carnitine (C_8_C) at 0.5 mg/g 20 min before treadmill running (A) or grip strength testing (B). A group of wildtype littermates were assessed as control. A “top speed” running protocol was used, which assesses the highest velocity at which the mice could complete a 5 min run (see Methods for details). C,D) The same experimental design as in Panels A and B, but using CrAT muscle-specific knockouts (CrAT^mKO^). *P<0.05; **P<0.01; ****P<0.0001 by Student’s T-test. All bars represent means and standard deviations.

## DISCUSSION

This study challenges the long-standing dogma of medium-chain triglyceride (MCT) metabolism and proposes a novel therapeutic application for acylcarnitines, specifically medium-chain acylcarnitines such as C_8_-carnitine, in the treatment of LC-FAODs. For decades, MCTs have been promoted as efficient energy sources that bypass the carnitine shuttle and rapidly fuel peripheral tissues. However, using both acute and chronic treatment paradigms in multiple mouse models, we demonstrate that MCT and triheptanoin fail to improve muscle function, exercise capacity, or metabolic stress in LC-FAODs. These findings reveal critical limitations of current therapies and motivate the exploration of alternative delivery strategies that effectively target the muscle and heart, such as acylcarnitines. Our data reposition acylcarnitines from their well-characterized role as mitochondrial transport intermediates and diagnostic biomarkers into potential metabolic therapeutics. Specifically, we show that orally administered medium-chain acylcarnitines are absorbed into the circulation intact, reach peripheral tissues, and are rapidly oxidized to provide energy. Notably, C_8_-carnitine not only supports oxidative metabolism but also releases free carnitine upon metabolism, helping to preserve the free CoA pool and thereby relieve mitochondrial stress. Further, whereas activation of free fatty acids such as C_7_ or C_8_ to their corresponding acyl-CoAs by ACSMs requires an input of ATP, conversion of acylcarnitines to acyl-CoAs requires no energy. These features offer a unique dual mechanism of action distinct from current therapies like triheptanoin.

A major mechanistic insight from our study is the identification of CrAT, rather than CPT2, as the critical enzyme mediating the oxidation of medium-chain acylcarnitines in heart and skeletal muscle. This challenges existing assumptions about the uniformity of carnitine shuttle function across acyl-chain lengths. While both CPT2 and CrAT can catalyze the conversion of medium-chain acylcarnitines to medium-chain acyl-CoAs *in vitro*, only CrAT appears to do so in intact mitochondria, as evidenced by the complete loss of C_8_-carnitine oxidation in CrAT-deficient muscle but no change in CPT2-deficient muscle. CPT2 has two small membrane-binding helices that are thought to mediate interaction with cardiolipin and CACT, the acylcarnitine translocase which is imbedded within the inner mitochondrial membrane [34]. Intriguingly, the CrAT three-dimensional structure has an alpha helix (helix 1) that aligns spatially with the membrane binding helices of CPT2 [34]. The physiochemical properties of this helix indicate it has a highly hydrophobic surface that would be predicted to bind membranes (Supplemental Figure 5). We speculate that CrAT may bind to the inner mitochondrial membrane and form a functional complex with CACT to facilitate the import and processing of medium-chain acylcarnitines, whereas CPT2 may fulfill this role for long-chain substrates.

Clinically, our findings suggest that medium-chain acylcarnitine therapy has broad and underexplored therapeutic potential. First, C_8_-carnitine robustly improves exercise performance and reduces markers of muscle damage (e.g., serum creatine kinase and lactate) in multiple preclinical models of LC-FAODs, including a new, more clinically faithful model of VLCAD deficiency. Unlike triheptanoin, C_8_-carnitine improves not only locomotion and grip strength but also metabolic homeostasis during physical exertion. This positions C_8_-carnitine as a candidate for pre-emptive, single-dose use in LC-FAOD patients prior to physical exertion, or as a chronic dietary supplement analogous to triheptanoin—but with potentially fewer gastrointestinal side effects due to its water-solubility and rapid clearance. Second, and notably, by demonstrating that medium-chain acylcarnitines can bypass CPT2 and utilize CrAT, we establish a mechanistic rationale for using C_8_-carnitine in CPT2-deficient patients.

Beyond LC-FAODs, medium-chain acylcarnitines may provide therapeutic value in a range of other metabolic disorders that have failed to respond to triheptanoin, such as McArdle’s disease, phosphofructokinase deficiency, GLUT1 deficiency, pyruvate carboxylase deficiency, and alternating hemiplegia of childhood [35-39]. Unlike MCT or triheptanoin, oral acylcarnitines require no lipases for processing and thus may be absorbed from the gut very quickly, allowing for energy supplementation in disorders where pancreatic insufficiency is common, such as cystic fibrosis. Our findings also carry implications for the broader field of sports nutrition. Despite the widespread use of MCT oil for weight loss and athletic performance, our data reveals that muscle and heart mitochondria are ill-equipped to directly metabolize medium-chain fatty acids due to a lack of ACSM expression. Most orally ingested MCT is sequestered by the liver, with little reaching peripheral tissues in usable form. In contrast, medium-chain acylcarnitines show greater systemic distribution and are efficiently utilized by muscle and heart. Importantly, long-term MCT use has been shown to increase liver triglyceride content [40]—a side effect unlikely to be shared by acylcarnitine formulations, which cannot be directly elongated or stored like medium-chain free fatty acids. Therefore, medium-chain acylcarnitines may offer a more targeted, effective, and safer supplement for exercise performance and recovery.

In summary, our study redefines the therapeutic landscape of fatty acid oxidation disorders by unveiling a previously unappreciated metabolic bottleneck—the lack of ACSM expression in target tissues—and by introducing medium-chain acylcarnitines as a compelling alternative to MCT-based therapies. We uncover CrAT as the critical mediator of medium-chain acylcarnitine metabolism in muscle and heart and provide preclinical evidence that oral C_8_-carnitine is bioavailable, well-tolerated, and effective. Future investigations should explore the pharmacokinetics, long-term safety, and dosing strategies of medium-chain acylcarnitines in humans. Structural studies to characterize CrAT–CACT interactions and tissue-specific delivery mechanisms will also be critical. Ultimately, this work opens new therapeutic avenues not only for LC-FAODs but also for a wide range of metabolic and neuromuscular disorders where energy failure is central to disease pathogenesis.

## METHODS

### Animals

All protocols for animal use and experiments were approved by the University of Pittsburgh Institutional Animal Care and Use Committee (IACUC), ensuring all work was done in accordance with the Animal Welfare Act and Public Health Service Policy on Humane Care and Use of Laboratory Animals. Mice and rats studied through this work were maintained in a pathogen-free barriered facility on a 12-hour light/dark cycle. LCADKO mice, generated by Dr. Philip Wood, were bred and genotyped as previously described [14]. The VLCAD^dKO^ strain was previously described [32]. CPT2 and CrAT conditional knockout strains were developed by Drs. Michael Wolfgang and Randall Mynatt, respectively, and have been well-described [27, 41-44]. To generate muscle-specific knockout strains, both were crossed with the Cg-Tg(ACTA1-cre)79Jme/L transgenic strain. All genotyping was performed by Transnetyx, Inc. (Memphis, TN).

### Acute and chronic lipid administration

Pelleted rodent diets containing either MCT (C_8_ triglyceride) or triheptanoin (C_7_ triglyceride) were manufactured by Research Diets, Inc. and were formulated to contain 35% of calories as MCT or triheptanoin, with an additional 10% of calories as long-chain fat to supply essential fatty acids. These diets were fed ad libitum for 10 days. For C_8_-carnitine feeding, C_8_-carnitine (ClearSynth, Brampton, ON) was mixed into powdered rodent diet at 3% w/w and fed in jars for 10 days. For acute administration of MCT, triheptanoin, and C_8_-carnitine, the lipids were either gavaged as the oils (MCT, triheptanoin) or dissolved in saline (C_8_-carnitine). For the latter, saline vehicle was given to the control animals. All dosing was 0.5 mg/g body weight, delivered 20 minutes prior to functional testing.

### Treadmill running

All treadmill experiments used an Ugo Basile treadmill (Gemonio, Italy). Mice were acclimated to the machine by walking at 5 m/min for 20 minutes for three consecutive days prior to exhaustion testing. On the day of testing, mice were fasted for 5 hr beginning at 7 a.m. Blood lactate and glucose were tested with handheld meters just prior to running. The running protocol was optimized for each strain. For LCADKO, the treadmill was programmed to accelerate from a starting pace of 5 m/min to 25 m/min over a 10 minute period; after reaching the maximum speed of 25 m/min they were run to exhaustion, defined as taking 25 cumulative foot shocks over the course of the exercise or remaining on the shock grid for > 5 seconds. For VLCAD^dKO^ mice, exercise intolerance was so severe that they could not reach a speed of 25 m/min. Therefore, the acceleration phase went from 5 m/min to a maximum of 18 m/min over 10 min. Exhaustion was defined using the same criteria as LCADKO. Muscle-specific CPT2 and CrAT knockout mice had much milder exercise intolerance than either LCADKO or VLCAD^dKO^. For these strains, a “top speed” running protocol was adapted from Petreyra et al [33]. After an initial walking pace at 5 m/min for 5 minutes, the mice were rested for 2 minutes; then, the speed was increased by increments of 5 m/min, with 2 min of rest in between each increment, until the mice reached a speed where they were unable to run for the full 5-minute trial, defined by the mouse taking 25 total foot shocks or remaining on the shock grid for more than 5 consecutive seconds. The “top speed” for each mouse was defined as the speed of the last successfully completed 5 min interval.

### Grip Strength Assessment

Mice were fasted for 5 hours prior to testing. Baseline measurements were taken using a BIOSEB GT3 grip strength test machine (BIOSEB, North Pinellas, FL) fitted with a metal gripping grid, measuring grams of pull. Each mouse was held by the tail while the front legs were allowed to grip the grid. The tail was then gently pulled until the mouse released its grip. Each mouse was measured 5 times to gather a representative number of measurements. The pull data were normalized to body weight.

### Open Field Actimeter Measurements

The BIOSEB Infrared Actimeter system (BIOSEB, North Pinellas, FL) utilizes an X-Y-Z axis infrared laser system to track maximum speed, mouse locomotion, total distance traveled in actimeter, and the percent of time resting in the actimeter. Mice were put into the open field actimeter system after a 5 hr fast and locomotion metrics were recorded by the BIOSEB system for 20 minutes.

### Creatine Kinase Activity Assay

For baseline CK, blood was collected early in the light cycle under fed conditions. Serum was collected and used with the Creatine Kinase Colorimetric Activity Assay Kit following the manufacturer’s instructions (Abcam, Boston, MA, AB155901). To induce rhabdomyolysis, we used a submaximal eccentric treadmill protocol [45]. Eccentric running was sustained at 12 meters/minute until exhaustion. Blood was sampled 24 hr later and used to measure CK as above.

### ACSM expression

First, we queried our previously published proteomics dataset [20] for ACSM proteins in wildtype, fed-state liver, quadriceps, and heart. The peak intensities for ACSM isoforms 1-5, from N=5 male 129S1 mice, were plotted as a heatmap in Graphpad Prism 10.0. This data was verified by immunoblotting. Fifty ug of total liver, muscle, or heart proteins were electrophoresed on Criterion SDS polyacrylamide gels (BioRad, Hercules, CA) and transferred to nitrocellulose membranes. Primary antibodies used were anti-GAPDH (Proteintech, 10494-1-AP, 1:1000 dilution), anti-ACSM1 (Invitrogen, PA5-32119, 1:1000), anti-ACSM2 (Abcam, ab181865, 1:500), anti-ACSM3 (Invitrogen, PA5-100374, 1:1000), and anti-ACSM5 (ProteinTech, 16591-1-AP, 1:1000). Membranes were incubated overnight at 4°C with primary antibodies, then incubated with HRP-conjugated secondary Goat Anti-Rabbit or Goat Anti-Mouse IgG antibody (Bio-Rad, 1706515 and OBT1500P, Hercules, CA) at 1:2000 dilution for 1-2 hours at room temperature. Clarity Max ECL kit (Bio-Rad, Hercules, CA) was used to visualize protein bands and images of the membranes were captured using the Protein Dock system (FluorChem M System, Bio-Techne, Minneapolis, MN).

### Tissue acylcarnitine analysis

LCADKO mice were fed either powdered standard rodent diet or the same diet mixed with C_8_-carnitine (3% w/w) for 10 days. All mice were sacrificed in the fed state, early in the light cycle. Liver, heart, and quadriceps tissue were snap frozen for acylcarnitine profiling in the Metabolic Core Facility at the University of Pittsburgh/UPMC Children’s Hospital of Pittsburgh as described [46, 47]. Frozen tissue (30 – 60 mg) was wrapped in aluminum foil and suspended in a liquid nitrogen bath for 2 min, then removed, placed on a hard benchtop and the tissue pulverized with a hammer. The pulverized tissue was transferred to a pre-weighed microcentrifuge tube (MCT) and a post-weight determined. An aliquot (20 μL) of isotope labeled carnitine standards (reconstituted in methanol) was spiked into the MCT along with 280 µL of ethanol (abs.), then vortex-mixed for 1 min to extract the acylcarnitines. The MCT was placed in a sonication ice water bath (4 – 22 °C) and sonicated for 30 min with vortex-mixing (30 sec every 10 min). After sonication, 700 µL of ethanol (abs.) was added to the MCT and vortex-mixed for 30 sec. The extracted sample was centrifuged (13,000xg, 10 min, 4°C). A portion of the supernatant (50 μl) was dried under a stream of nitrogen gas and the acylcarnitine butyl esters generated by reaction (60°C for 15 min) in 100 μl of 3 N HCl in butanol. Dried residues were reconstituted in acetonitrile– water (80:20) for flow-injection ESI-MS–MS analysis. Analysis was performed on a triple quadrupole API4000 mass spectrometer (AB Sciex™, Framingham, MA) equipped with an ExionLC™ 100 HPLC system (Shimadzu Scientific Instruments™, Columbia, MD). Acylcarnitine standards were purchased from Amsterdam UMC—VUmc (Amsterdam, NL) and Cambridge Isotope Laboratories, Inc. (Andover, MA). Acylcarnitines were measured using multiple reaction monitoring (MRM) for free carnitine (C_0_, m/z 218 ⇒ m/z 103) and acetylcarnitine (C_2_, m/z 260 ⇒ m/z 85) and precursor scan for precursor ions (Q1) of acylcarnitines (C_3_ to C_18_, scan range m/z 270 to 502) that generated a product ion (Q3) at m/z 85.

### Mitochondrial isolation

For Oroboros respirometry, fresh mitochondria were isolated from mouse skeletal muscle, heart, and liver using differential centrifugation. All protocols for mitochondrial isolation were adapted from MitoPedia, the official Oroboros reference site (https://wiki.oroboros.at/index.php). After tissues were excised from the mice, all steps were carried out on ice or in 4°C conditions. Conditions varied slightly for each tissue type as follows. <u>Skeletal muscle:</u> tissue was placed in ice cold phosphate buffered saline (PBS) containing 10 mM EDTA, minced and incubated in PBS with 10 mM EDTA and 0.05% Trypsin for 30 minutes on ice to permeabilize the tissue. Tissue was centrifuged at 200 x g to remove the trypsin buffer and replaced with IBM1 buffer containing 50 mM KCl, 67 mM sucrose, 50 mM Tris pH7.4, 10 mM EDTA, and 0.2% BSA before homogenization via mechanical shearing using a glass Dounce homogenizer. The homogenate mix was centrifuged at 700 x g for 10 minutes to remove cellular and tissue debris. Then, the mitochondria-containing supernatant was transferred to a new tube and centrifuged at 8,000 x g for 10 minutes. The mitochondrial pellet was resuspended in IBM2 buffer containing 250 mM sucrose, 3 mM EGTA and 10 mM Tris pH 7.4 before a final 8,000 x g centrifugation for 10 minutes. Resulting intact mitochondria were resuspended in Elution Buffer containing 250 mM sucrose, 0.02 mM EGTA, 10 mM Tris pH 7.4, 2 mM Tris-phosphate pH 7.4, and 5 mM MgCl2. <u>Heart</u>: freshly isolated hearts were washed three times with ice cold PBS to remove excess blood before being placed in Buffer A containing 180 mM KCl, 4 mM EDTA, and 10 mg of BSA. Heart tissue was then minced and rinsed with ice cold Buffer B containing 180 mM KCl, 4 mM EDTA, 10 mg of BSA, and 2.5 mg subtilisin to rinse off any remaining Buffer A before being incubated on ice in Buffer B for 10 minutes to allow the subtilisin to act as a detergent. After 5 minutes of centrifuging at 200 x g, minced heart tissue was resuspended in Buffer A and homogenized by mechanical shearing. Cell and tissue debris were pelleted by 1000 x g centrifugation for 10 minutes, after which the supernatant was transferred to a fresh tube and centrifuged at 6,200 xg for 10 minutes. Intact mitochondria were resuspended in Buffer C containing 180 mM KCl, and 4 mM EDTA. <u>Liver</u>: fresh tissue was excised into ice-cold buffer containing 225 mM mannitol, 75 mM sucrose, and 0.2 mM EDTA. The liver was then minced and homogenized by mechanical shearing before a 10-minute centrifugation at 1,000 x g. The supernatant was transferred to a fresh tube and centrifuged for 10 minutes at 6,200 x g. Intact mitochondria were resuspended in Buffer C, described above, centrifuged once more at 6,200 xg for 10 minutes to wash the mitochondria, and finally resuspended in Buffer C, described above.

### Mitochondrial respirometry

An Oroboros Oxygraph-2K (Oroboros, Österreich, Austria) was used to assess the level of respiration by freshly isolated mitochondria (described above). 30 µL of liver, heart or skeletal muscle mitochondria was added to O2-equilibrated chambers containing 2 mL of MiR05 buffer (0.5 mM EGTA, 3 mM MgCl2, 60 mM lactobionic acid, 20 mM taurine, 10 mM potassium diphosphate, 20 mM HEPES, 110 mM D-sucrose). Once the oxygen concentration and change in oxygen concentration became stable at baseline, 10 µM of cytochrome C (Sigma, St. Louis, MO) was added to both chambers to assess mitochondrial membrane integrity. Subsequently, 5 mM malate (Sigma, St. Louis, MO) was added to both chambers to support oxaloacetate production, fueling the TCA cycle. 2 mM ADP was then added to support ATP production from the electron transport chain. Finally, either 50 µM of medium-chain free acids or acylcarnitines were added and maximal respiration response recorded. An N=4 or more was performed for all comparisons. All Oxygraph-2K signals were normalized to mitochondrial protein content as determined by BCA protein assay kit (Thermo Fisher Scientific, Waltham, MA).

### Lipid oral bioavailability

Sprague Dawley Rats fitted with externalized jugular access ports were purchased from Charles River (Wilmington, MA). The experimental design was to dose with 0.5 mg/g of either triheptanoin or C_8_-carnitine, comparing blood concentrations over time following i.v. doses to those of oral doses. For oral dosing, rats were briefly incapacitated using isoflurane and then gavaged with 0.5 mg/g of body weight of either triheptanoin or C_8_-carnitine via oral gavage. The rats quickly regained consciousness. Blood samples were drawn from the jugular IV port of each awake, dosed rat at t=0 minutes, 10 minutes, 20 minutes, 30 minutes, 60 minutes, 90 minutes, and 120 minutes. For i.v. dosing, triheptanoin oil was not used as i.v. administration of oils is toxic. Rather, an equal amount of C_7_ free fatty acid was dissolved in 0.9% saline solution and injected into the port of awake rats. C_8_-carnitine was administered in the same manner. All blood samples were centrifuged, and serum was snap-frozen and stored at -80°C until time of analysis by mass spectrometry. Serum samples were analyzed via tandem mass spectrometry (MS/MS) for free fatty acid quantification by the Mass Spectrometry Core at the University of Pittsburgh, or for serum acylcarnitine profiling by the Metabolic Core at the University of Pittsburgh/UPMC Children’s Hospital of Pittsburgh. Prior to C_7_ analysis, the serum lipids were hydrolyzed to release any C_7_ from intact triglycerides.

### Acyl-CoA synthetase activity

Initial assays with C_8_ followed the conversion of ^14^C-C_8_ into ^14^C-C_8_-CoA as we previously described [40]. Briefly, mitochondria were isolated by differential centrifugation and resuspended in cold SET buffer (10 mM Tris–HCl, 250 mM sucrose, 1 mM EDTA). Synthetase reactions (200 µl) contained 5 µl of homogenate with 10 µM ^14^C-fatty acid in a buffer of 40 mM Tris–HCl, 5 mM ATP, 5 mM MgCl_2_, 4 mM CoA, 0.8 mg/ml Triton WR1339, and 1 unit/ml inorganic pyrophosphatase. After 2 min incubation at 37 °C, reactions were stopped with sulfuric acid and extracted four times with ether prior to scintillation counting. Data were normalized to protein concentration. Later experiments used tandem mass spectrometry to directly detect the acyl-CoAs formed in the reactions. Freshly prepared tissue homogenates (250 µg) were incubated with 50 µM of either free fatty acid (C_7_, C_8_, C_16_) or the acylcarnitine conjugates of these fatty acids, in a reaction buffer containing 40 mM Tris (pH=8), 5 mM ATP, 5 mM MgCl_2_, and 1 U/mL inorganic pyrophosphatase. The reactions were started by addition of free CoA to a final concentration of 200 µM. Reaction tubes were placed in a water bath shaker at 37°C for one hour. Reactions were then flash frozen in liquid nitrogen and stored at -80°C. The medium-chain CoA concentration of each reaction tube was measured via tandem MS/MS analysis at the Biochemical Genetics Laboratory at Colorado’s Children’s Hospital (Aurora, CO).

### Carnitine acyltransferase expression, purification, and activity

Expression constructs for human CrAT (pJH15-pET21-CRAT-6His, gift from Dr. Chaitan Khosla, Stanford) and human CPT2 (pET21b-CPT2-6His) were transformed into *E coli* and expressed overnight at 18°C, by addition of 0.5 mM IPTG. Bacterial pellets were lysed via sonication and the lysates clarified by ultracentrifugation. Supernatants were applied to a preequilibrated ion-exchange chromatographic column HisTrap HP on an ӒKTA pure™ chromatography system (Cytiva, USA). The column was washed 20 column volumes of washing buffer (50 mM phosphate, 10 mM imidazole, 300 mM NaCl, 10% glycerol, pH 8). Recombinant protein was eluted by increasing the percentage of elution buffer (50 mM phosphate, 250 mM imidazole, 300 mM NaCl, 10% glycerol, pH 8). Eluted pure CrAT and CPT2 proteins were dialyzed against above phosphate buffer without any imidazole and then concentrated and kept frozen in small aliquots at -80°C. Enzyme activity in the direction of acylcarnitine → acyl-CoA was measured with a novel coupled assay using the fluorescence of electron transferring flavoprotein (ETF) as the readout. In this assay, illustrated in Supplemental Fig 6, recombinant CrAT or CPT2 were incubated with C_7_-carnitine or C_8_-carnitine to generate C_7_-CoA or C_8_-CoA, which were then dehydrogenated by recombinant medium-chain acyl-CoA dehydrogenase (MCAD), which passes electrons to its natural electron acceptor ETF. ETF is a fluorescent protein which becomes quenched upon accepting electrons. We have previously published a method using recombinant ETF to follow the activity of acyl-CoA dehydrogenases [48]. Here, each 200 µl reaction contained 50 mM Tris (pH8), 0.5% glucose, glucose oxidase (∼20U/ml), catalase (∼10U/ml), 2 µM recombinant ETF, 1 µg recombinant MCAD, 0.2 µg recombinant CPT2 or CrAT (MCAD/CPT2 or MCAD/CrAT ratios were 5:1), and 0.5 mM CoA. After incubation at 32° for 2 minutes, baseline fluorescence was recorded for 1 minute at Ex 340 nm / Em 490 nm on a FLUOstar Omega plate reader (BMG Labtech). Then, the reaction was started by addition of C_7_-carnitine or C_8_-carnitine to a final concentration 25 µM and the fluorescence intensity was recorded for 1 minute. The slope and Y-intercept were used to calculate activity as we have previously described [48].

### Statistical Analyses

All statistical analyses were performed in GraphPad Prism (version 10.3, Graphpad Prism Software Inc., San Diego, CA). Two-tailed student’s t-tests were used to compare treated to untreated, knockout to wildtype, and other two-group comparisons. An area under the curve (AUC) calculation was used to determine the oral and IV bioavailability of triheptanoin and C_8_-carnitine. After constructing a ratio of oral AUC/IV AUC, a t-test was used to compare the calculated overall bioavailability. Oroboros oxygraph data were generated in DatLab (version 8.0, Oroboros Instruments, Innsbruck, Austria) and exported to Microsoft Excel where the average peak difference in oxygen consumption was calculated. All data were subsequently transferred to Prism for student’s t-test analyses.

## Acknowledgements

Acylcarnitine profiling was performed in the Metabolic Core Facility (RRID:SCR_025224) located within the John G. Rangos Research Center, UPMC Children’s Hospital of Pittsburgh and services and instruments used in this project were graciously supported, in part, by the University of Pittsburgh, Department of Pediatrics.

