## Supplemental figures for "Oral octanoylcarnitine alleviates exercise intolerance in mouse models of long-chain fatty acid oxidation disorders"

Figure S1

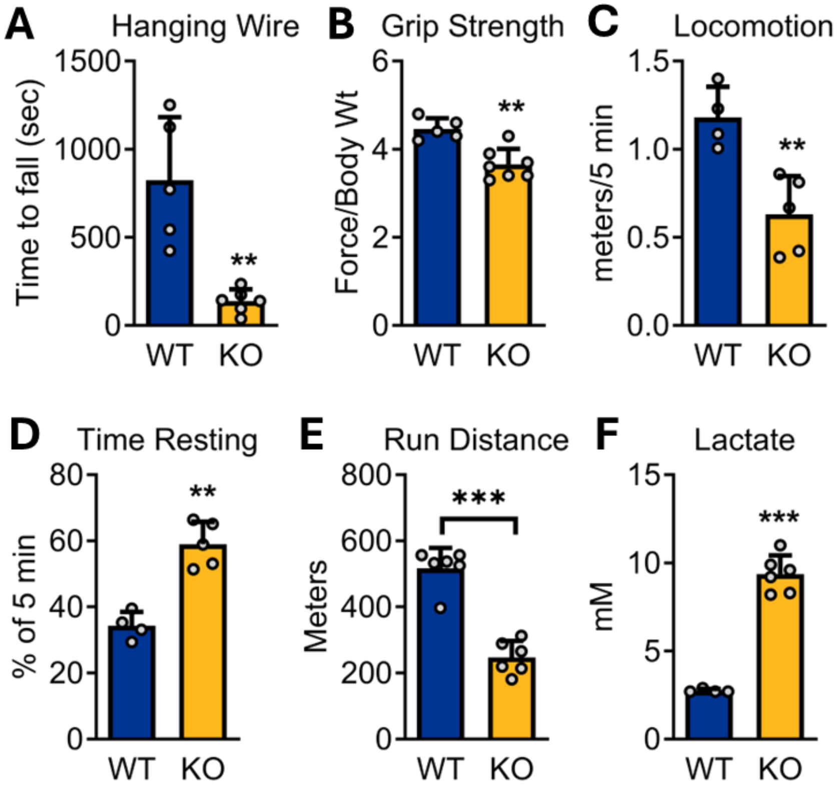


**Supplemental Figure S1**. Muscle function in LCADKO (KO) and wildtype (WT) control mice. A) Hanging wire test: WT and KO mice were suspended from a wire 50 cm above the surface. Shown is the time to fall. B) Forelimb grip strength measured with a force meter and normalized to body weight. C,D) An open field actimeter was used to record total basal movement (C) and percentage of time spent resting over a 5 minute interval. E,F) Mice were challenged with acute treadmill run to exhaustion protocol. Shown are the total distance run and the post-run blood lactate. **$P<0.01$; ***$P<0.001$; by Student's T-test. All bars represent means and standard deviations.

Figure S2

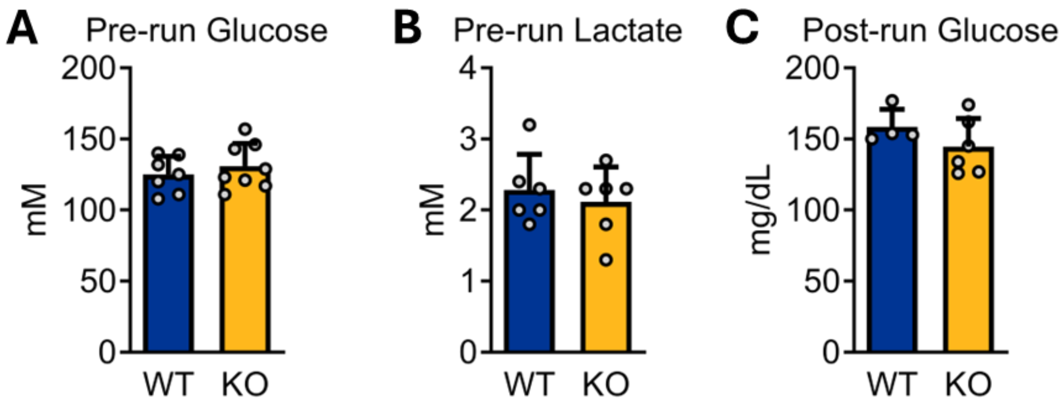


**Supplemental Figure S2**. Blood glucose and lactate in LCADKO (KO) and wildtype (WT) control mice. A) Blood glucose just before treadmill challenge, and B) blood lactate just before treadmill challenge. C) Glucose after exhaustion. All bars represent means and standard deviations.

Figure S3

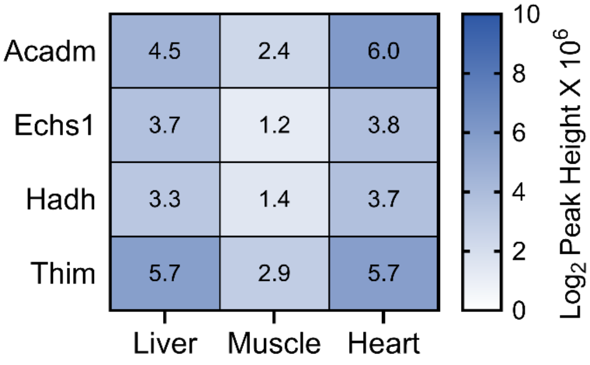


**Supplemental Figure S3**. Protein expression of medium-chain mitochondrial FAO enzymes, taken from a previously published mass spectrometry proteomics dataset [20]. Shown are $\log_2$ peak heights X $10^6$ for enzymes catalyzing steps 1-4 of medium-chain β-oxidation: 1) medium-chain acyl-CoA dehydrogenase (Acadm); 2) mitochondrial enoyl-CoA hydratase (Echs1); hydroxyaycyl-CoA dehydrogenase (Hadh); and mitochondrial 3-ketoacyl-CoA thiolase (Thim).

Figure S4

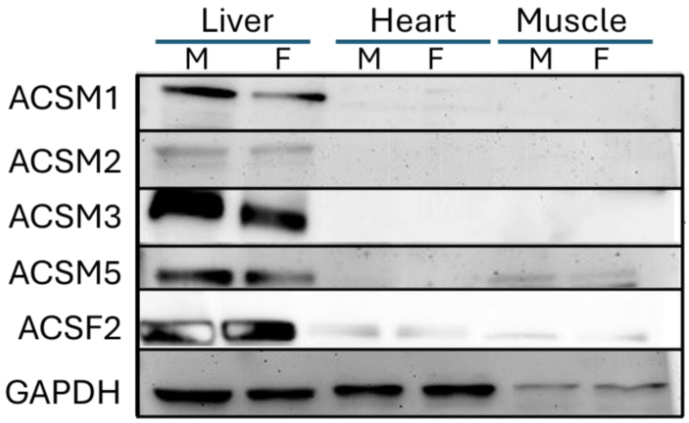


**Supplemental Figure S4**. Immunoblotting of putative medium-chain acyl-CoA synthetases (ACSMs) in wildtype male (M) and female (F) mouse liver, heart, and muscle. There are five ACSM family members (ACSM1 to ACSM5). No antibody was commercially available for ACSM4, which is a gut-specific enzyme. In addition to the ACSMs, acyl-CoA synthetase family member-2 (ACSF2) has been shown to have activity with medium-chain fatty acids *in vitro* and thus was included here. GAPDH was the loading control.

Figure S5

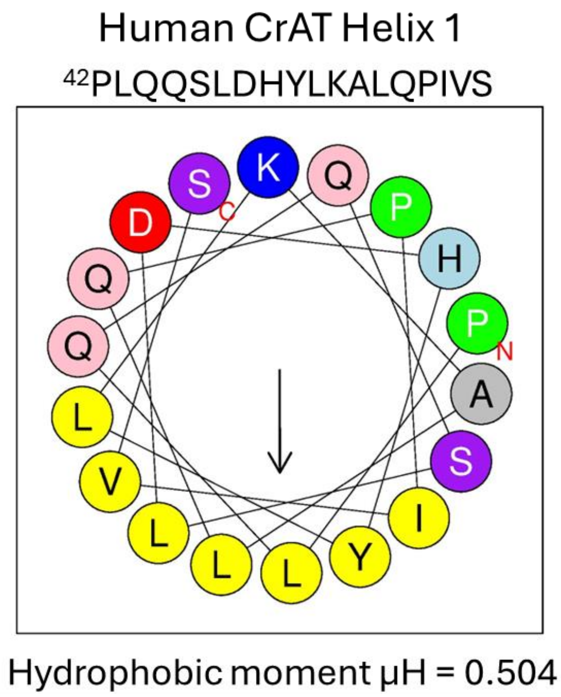


**Supplemental Figure S5**. Predicted membrane-binding helix in human CrAT. Helix 1 of CrAT (residues 42-59) has a strong hydrophobic moment with a hydrophobic face (indicated in yellow), which is characteristic of membrane-associated helices. Analysis performed in Heliquest (https://heliquest.ipmc.cnrs.fr).

Figure S6

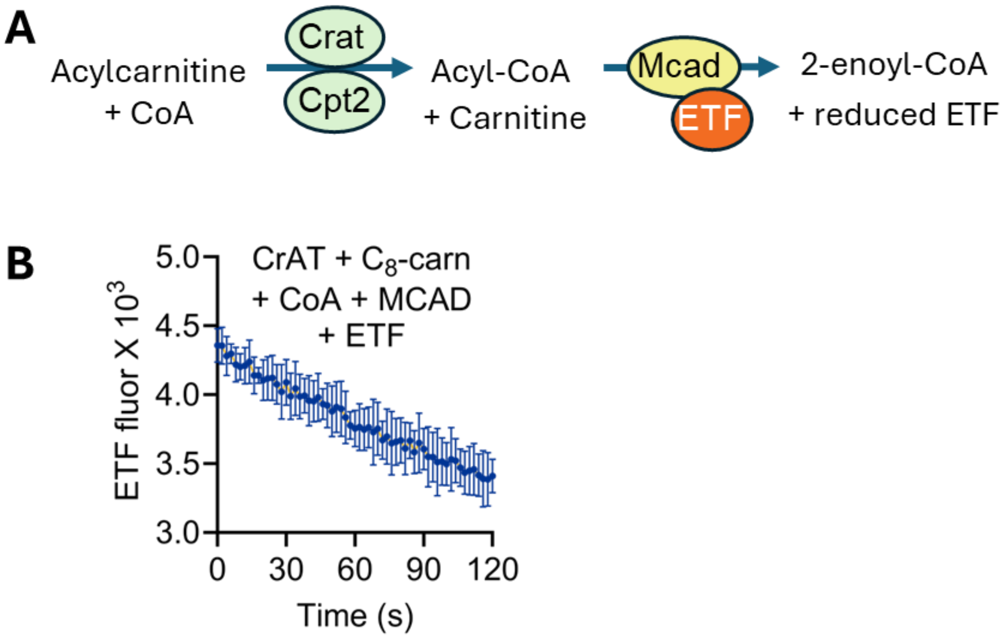


**Supplemental Figure S6**. Coupled acylcarnitine transferase activity assay with CrAT/CPT2, MCAD and ETF. A) Scheme of assay. Recombinant CrAT or CPT2 converts acylcarnitine substrate and CoA into acyl-CoA. Recombinant MCAD dehydrogenates the medium-chain acyl-CoA and passes electrons to electron-transferring flavoprotein (ETF). B) Proof-of-concept for the assay: shown is the reduction of ETF (loss of fluorescence) over time during reaction between CrAT, $C_8$-carnitine, free CoA, and MCAD. The reaction was performed in triplicate and the graph shows mean and standard deviation. The slope and Y-intercept can be used to enzyme calculate activity.
